# Neural underpinnings of action adaptation in the subthalamic nucleus

**DOI:** 10.1101/2022.06.28.497904

**Authors:** Damian M. Herz, Manuel Bange, Gabriel Gonzalez-Escamilla, Miriam Auer, Muthuraman Muthuraman, Rafal Bogacz, Sergiu Groppa, Peter Brown

## Abstract

Adapting our actions to changing goals and environments is central to intelligent behavior. There is substantial evidence that the basal ganglia play a crucial role in reinforcing actions that have led to favorable outcomes. However, little is known about the neural mechanisms underlying action adaptation following unfavorable outcomes when change is warranted. Here, we recorded electrophysiological activity and applied bursts of electrical stimulation to the subthalamic nucleus (STN), a core area of the basal ganglia, in patients with Parkinson’s disease using deep brain stimulation electrodes. During a task where patients continuously had to adapt their force depending on changing action-value associations, decreases in STN beta (13-30 Hz) activity in two critical time windows were associated with poorer outcomes and stronger action adaptation. STN stimulation reduced beta activity and led to stronger action adaptation if applied within the time windows when STN activity reflected action evaluation and adaptation. These results suggest that dynamic modulation of STN activity facilitates adaptive behavior.

## Introduction

To successfully navigate between affordances offered by our environment we have to learn how actions differ regarding their usefulness and update these associations if they no longer lead to desirable outcomes ^1-3^. This does not only apply to deciding what action to choose, but also how to perform it. For example, during foraging agents do not only need to choose between e.g. eating grapes or nuts depending on their nutritional value, but also learn how much force to apply for cracking a nut without crushing it. In the reinforcement-learning framework, action-value associations are learned by comparing actual and expected value for different options and then adapting actions accordingly ^3-5^.

Neurobiologically, there is strong evidence that firing rates of dopaminergic midbrain neurons and consequent striatal dopamine release increase when the actual value surpasses the expected value ^6-9^. With the coincident presence of glutamate in striatal synapses, this signal is thought to increase excitability of the respective cortical neurons and induce plasticity mechanisms through the cortico-basal ganglia direct pathway ^3, 10-14^. Thus, neural activity patterns leading to successful outcomes will be strengthened making the action more likely in the future. However, when actions are unsuccessful changes in movement are warranted and how this is achieved remains less clear.

Patients with Parkinson’s disease (PD) undergoing deep brain stimulation (DBS) offer the unique opportunity to directly record electrophysiological activity from the subthalamic nucleus (STN) in humans. The STN is a core part of the indirect (and hyperdirect) basal ganglia pathway exerting a net inhibitory effect on thalamo-cortical connections. Previous STN recordings in PD patients have shown that movement-related activity in the beta-band (∼13-30 Hz) is strongly modulated during force production ^15-18^. Importantly, beta activity is reduced when force is adjusted irrespective of whether this entails an increase or a decrease in force ^19^. In other words, decreases in beta activity appear to be reflective of changes in force rather than force per se. A relationship between STN beta activity and concomitant movement changes has also been observed after the movement is terminated. For example, it has been demonstrated that movement errors due to external perturbations reduce STN beta activity compared to correct movements, but only if this error is relevant for action adaptation ^20^. Another study found that STN beta power after the movement is relatively higher when an action outcome has been favorable and the action should be repeated, and relatively lower after suboptimal actions necessitating change ^21^.

However, in these previous studies it was already obvious to the participants how the movement should be adjusted at the time when they observed a difference between actual and expected outcome conflating action evaluation and preparation. In other words, the observed changes might primarily have reflected movement preparation of the next movement rather than evaluation of the previous movement. Furthermore, it remains elusive whether these STN activity changes are merely correlative in nature or if they causally contribute to action evaluation and adaptation. This is particularly important since current approaches of therapeutic adaptive DBS employ STN beta activity as a feedback signal, which in turn is modulated by stimulation ^22, 23^.

To address these outstanding issues, we conducted electrophysiological recordings and applied bursts of electrical STN stimulation in 16 PD patients who had undergone DBS surgery. Based on a previous study of decision-making under uncertainty ^24^ we designed a force adaptation task in which participants continuously had to adapt their grip force based on the value associated with their previous action reflecting how close the actual force was to the target force. Importantly, the Value-feedback was dissociated from a second feedback cue necessary for action adaptation, so that patients could not infer with certainty how to adapt their force until after the second cue. We hypothesized that STN beta activity would be modulated by action-value feedback and consequent adaptation of grip force. Moreover, we expected extrinsic modulation of beta activity through direct electrical stimulation to modify trial-to-trial action adaptation.

## Results

During the task participants attempted to produce forces as close as possible to a target force in order to collect a maximum number of points. While they were aware of the approximate target force level on the first trial (∼20-25% of their maximum voluntary contraction, MVC), the target force changed over trials without being explicitly shown on the screen so that participants had to infer target force based on the feedback they received after each movement. The first feedback cue (Value-feedback) indicated how close the actual force was to the target force (ranging from 0 (worst) to 10 (best)) and the second feedback cue (Direction-feedback) showed whether the force had been too low or too high (figure 1A). Thus, after the Value-feedback participants could only infer how much change in force was necessary on the next trial, but not how this should be implemented (increase or decrease in force). Target force levels varied according to a noisy Gaussian decaying random walk with a mean of ∼20% MVC (figure 1B). Thus, throughout the task participants had to evaluate the value of their previous actions and adapt their force levels on the next trial accordingly.

**Figure 1.**
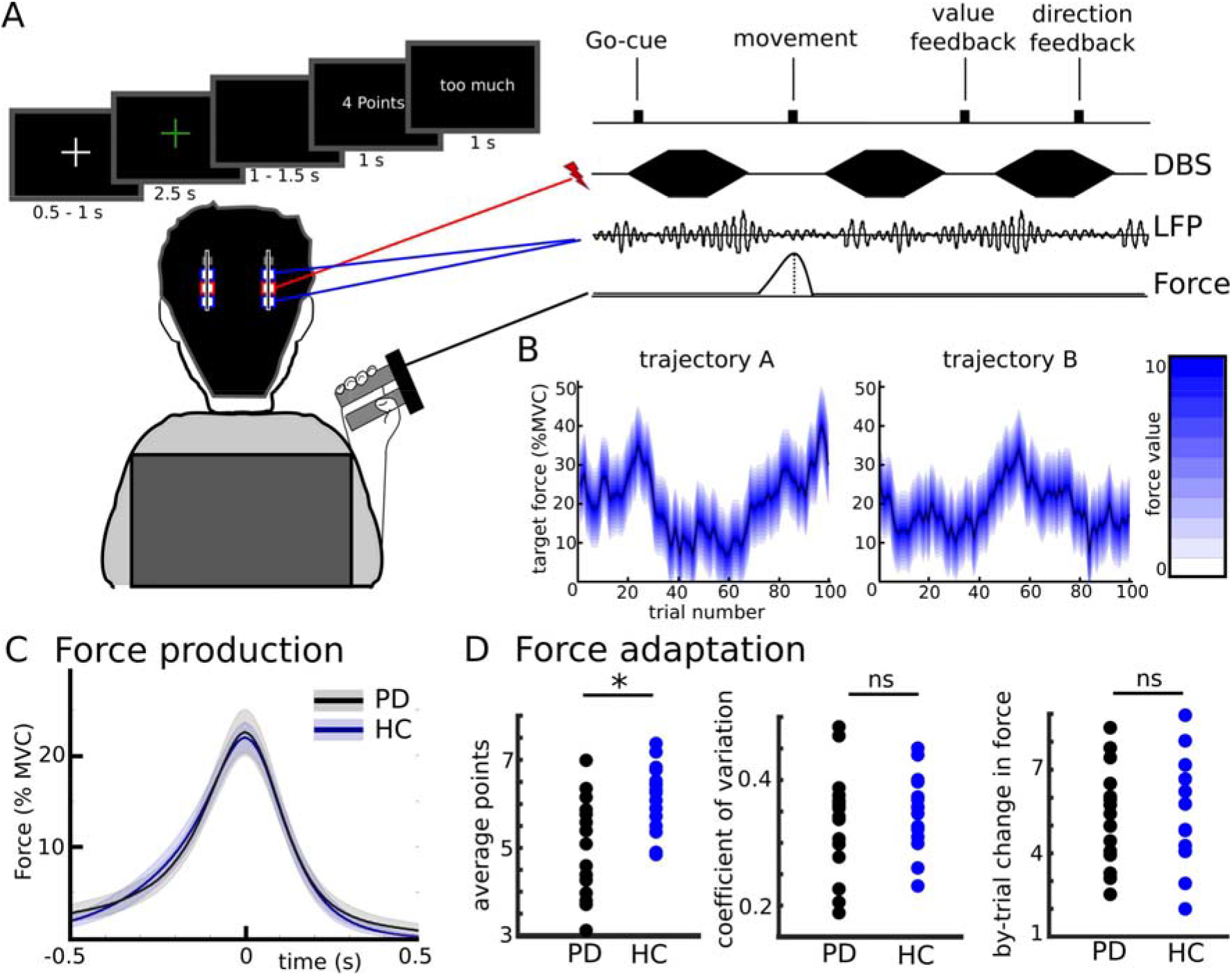
Paradigm and behavioral results. **A.** After the Go-cue (green fixation cross) participants exerted a certain force to match a target force which had to be inferred based on feedback regarding distance between actual and target force (Value-cue, here 4 points) and direction (had the previous force been too little or too much). Local field potentials (LFP) were recorded from bilateral subthalamic nucleus in two sessions, together with bursts of deep brain stimulation (DBS) in the second session **B.** The target force varied according to a Gaussian decaying random walk. Since patients participated in two sessions, two trajectories were used. **C.** Mean force production of patients and healthy controls. Shaded regions represent S.E.M. **D.** Single subject values of different force adaptation measures. PD, Parkinson’s disease; HC, healthy controls; MVC, maximum voluntary contraction; ns, not significant; * indicates a significant difference.

### Force production and adaptation in PD patients and healthy controls

Before analyzing the neural correlates of Value-feedback evaluation and action adaptation in the STN we assessed to what extent the performance of PD patients (n=16) was comparable to that of healthy people (n=15). In a first step, we investigated whether PD patients were able to produce forces similarly to healthy controls (HC). We compared multiple measures of force production including the MVC, mean peak force and its temporal derivative (yank), absolute force exerted and reaction time. None of these measures differed between groups (mean force trace shown in figure 1C, all statistics are listed in suppl. table 2) showing that patients can express normal force grips when force levels are relatively low. Next, we tested whether participants were able to adapt their actions according to the feedback they received. We found a strong correlation between actual and target force (average rho = 0.486) as well as between the Value-feedback and corresponding absolute change in force on the next trial (average rho = −0.419) in both groups (all P-values < 0.001) showing that they were able to follow task instructions and adapt their force according to the feedback. Patients showed overall poorer task performance compared to healthy controls regarding the difference between actual and target force resulting in lower average Value-feedback (t_29_ = 3.416, d = 1.23, P = 0.002, see figure 1C, suppl. table 2 and suppl. text). However, these difficulties did not impact patients’ overall ability for force adaptation as indicated by similar force level variability (coefficient of variation, t_29_ = 0.572, d = 0.21, P = 0.572) and mean by-trial absolute change in force (t_29_ = 0.346, d = 0.12, P = 0.732) between groups (figure 1D). Thus, the main interest of the current study, i.e. how actions are linked to values and how this is used for action adaptation, were similar in PD patients and healthy people.

### Modulation of STN beta power reflects action evaluation and adaptation

During the task we recorded local field potentials (LFP) directly from the STN through temporarily externalized DBS electrodes in PD patients. First, we asked whether modulations of STN beta activity reflected information contained in the two feedback cues. Aligning STN beta power to the feedback showed a strong (∼15%) decrease after the Value-feedback while changes after the Direction-feedback were minimal (figure 2A). To assess whether this reflected task-relevant information we conducted single trial regression analyses using Value-feedback (ranging from 0 to 10) and Direction-feedback (ranging from − 10, i.e., > 10% MVC too little force, to +10, i.e., >10% MVC too much force) as predictors and STN beta power as dependent variable using a sliding window approach (see methods for details). This revealed a significant, positive relationship between STN beta power and Value from 400-700 ms after the Value-cue (P_cluster_ < 0.05, figure 2B) while there was no significant relationship with Direction (figure 2C). Thus, STN activity during feedback reflected the absolute difference in force (i.e. Value-feedback) with lower STN beta power being related to lower Value-feedback, but not whether the force had been too high or too low.

**Figure 2.**
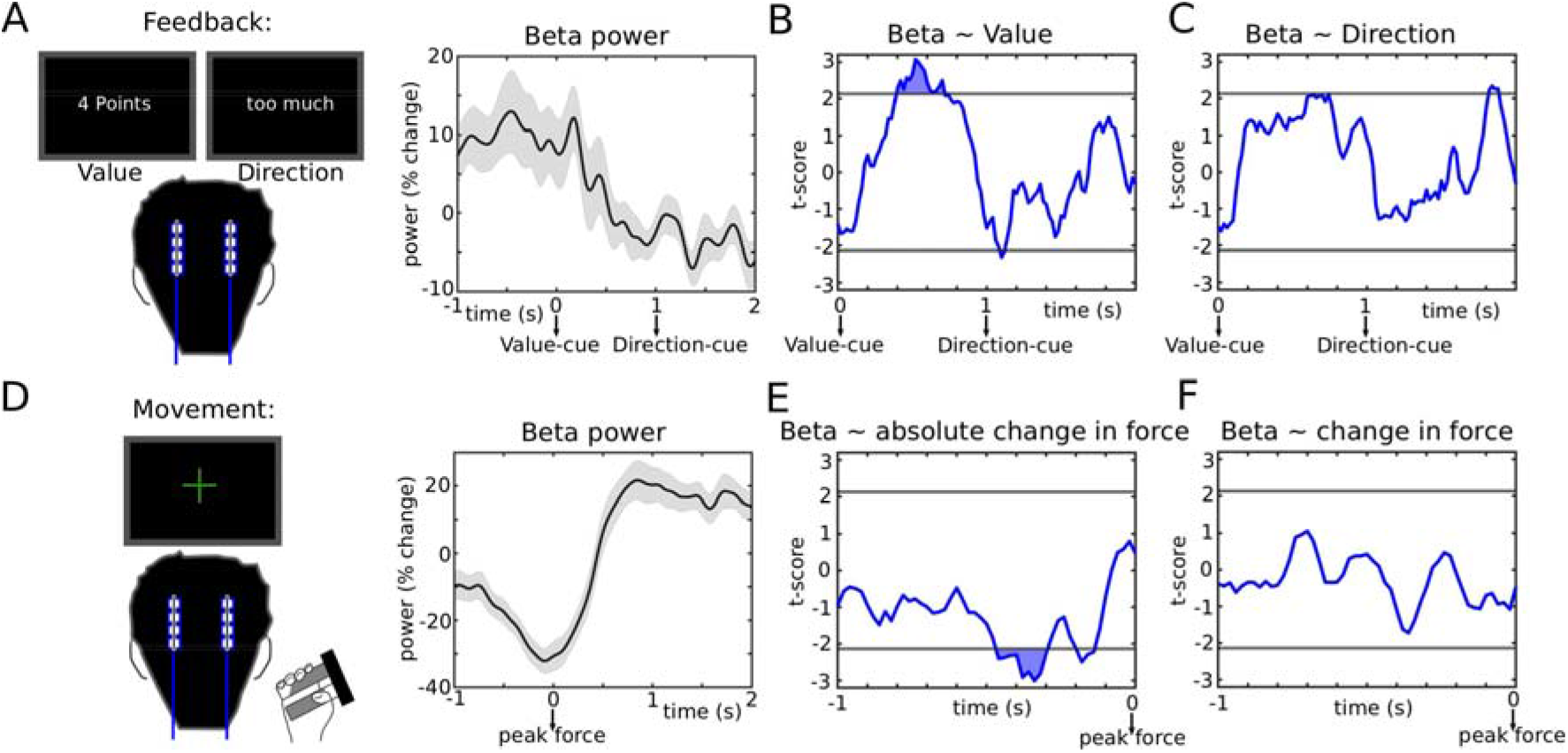
Correlates of action evaluation and adaptation. **A.** Subthalamic nucleus (STN) activity during the feedback period. STN beta power (∼13-30 Hz) only decreased after the Value-feedback. **B.** Regression between single trial measures of Value-feedback (ranging from 0 to 10) and beta power showing a significant relationship in a time window from 400 to 700 ms after the Value-cue. **C.** Same as B for regression between Direction-feedback (ranging from −10 to +10) and beta power. **D.** STN activity during the movement. STN beta power decreased strongly during the movement reaching the trough roughly at the peak force. **E.** Regression between single trial measures of absolute change in force and beta power showing a significant relationship in a time window from 460 to 300 ms before peak-force. **F.** Same as B for regression between change in force (positive values if force was higher, negative values if force was lower compared to previous trial) and beta power. In B, C, E and F horizonal grey lines show the cluster-building threshold and filled areas indicate significant clusters. Shaded areas in B and D represent S.E.M.

Since we also found a strong increase in STN alpha power in a short time window 0-500 ms after the Value-feedback in the time-frequency spectrum (suppl. figure 2A) we conducted an additional regression analysis using mean alpha power from this time window as dependent variable. However, there was no significant relationship with Value (t = 0.726, P = 0.468) or Direction (t = 0.948, P = 0.343).

Next, we asked whether modulation of STN beta activity also reflected action adaptation. To this end, we aligned STN beta power to the movement, which revealed a strong (∼30%) decrease in beta power reaching its trough around the peak force (figure 2D). We then conducted the same sliding-window regression analysis as above, now using absolute change in force (lower bound at 0 corresponding to no change) and change in force (positive values indicating an increase in force, negative values a decrease in force) as predictors. We found a significant, negative relationship between STN beta power and absolute change in force from 460-300 ms before peak force (P_cluster_ < 0.05, figure 2E). The lower beta power the stronger the absolute change in force. There was no significant relationship with change in force (figure 2F), i.e., STN beta power reflected how much participants adapted their force, but not whether it increased or decreased.

There was also a strong increase in STN gamma power in a short time window from −300ms until peak force (suppl. figure 2B). Therefore, we conducted an additional regression analysis using mean gamma power from this time window as dependent variable, which did not reveal a significant relationship between gamma power and absolute change in force (t=-0.760, P=0.448) or change in force (t=0.079, P=0.937).

Since Value-feedback and action adaptation were correlated (lower Value-feedback resulted in stronger absolute changes in force), it could be that STN beta power after the Value-cue was primarily related to action adaptation and that this might drive the correlation with Value-feedback. However, a control regression analysis between absolute change in force and STN beta power 400-700ms after the Value-cue (see above) did not show a significant effect (t=-0.357, P=0.721). While this suggests that STN beta power during the feedback was most closely related to the content of this feedback it does not necessarily entail that it was not relevant for action adaptation since the Value-feedback was informative of the necessary behavioral adaptations. To directly test whether STN activity played a causal role in action adaptation, participants performed the same task in a second session, where bursts of electrical stimulation were applied to the STN. Short bursts (mean duration: 250 ms) were given randomly throughout the task so that in any given 100 ms time window stimulation was applied in ∼50% of trials (suppl. figure 3A&B). This allowed us to compare timing-specific effects of STN stimulation on action adaptation without having to focus on any a-priori defined time windows. Based on the LFP regression analyses we hypothesized that STN stimulation during the feedback and movement period should affect how much participants adapted their force.

### STN causally contributes to action adaptation

First, we aligned the data to the feedback. Comparing trials in which stimulation had been applied to trials without stimulation in a sliding-window approach (see methods for more details) we found a significant, positive effect of STN stimulation on absolute change in force in a restricted time window from 180 to 340 ms after the Value-feedback (P_cluster_ < 0.05, figure 3B). Here, stimulation led to a stronger absolute change in force on the next trial. This did not depend on whether the force increased or decreased (suppl. figure 3C). Notably, this time period (hereafter termed DBS_value_) immediately preceded the window in which STN beta power reflected Value-feedback (400-700ms after the Value-cue, see figure 2B). Since previous studies have demonstrated that STN stimulation reduces beta power ^25-27^ this suggests that stimulation during DBS_value_ might have modulated STN beta activity in this time window. To test this, we analyzed STN beta power from the contacts surrounding the stimulation electrode during the stimulation session. Using common-mode rejection and artifact correction (see methods) we were able to recover the normal (i.e. as observed in the off stimulation session) feedback-modulation of STN beta power (see figure 3C & suppl. figure 4). As expected from previous studies ^25-27^ aligning STN beta power to onset of stimulation bursts showed a marked (∼40%) reduction in beta power (suppl. figure 5). We then extracted beta power from a time window of interest spanning DBS_value_ and the time window where STN beta power reflected Value-feedback (i.e. from 180 to 700 ms after the Value-cue, see grey rectangle in figure 3C) and compared it to control trials. In these control trials, stimulation also affected participants’ behavior but in a different time window (DBS_move_, see below). This analysis showed a significant reduction of beta power by stimulation during DBS_value_ from 280 to 500 ms after the Value-cue (P_cluster_ < 0.05, figure 3D, see black trace for beta power off stimulation). Thus, stimulation reduced STN beta power after the Value-cue, where beta activity normally reflected the absolute difference in force (i.e. Value-feedback) with lower beta power being related to larger differences necessitating stronger adaptation. Behaviorally, stimulation during DBS_value_ led to an increase in action adaptation as would be expected from trials with lower Value-feedback.

**Figure 3.**
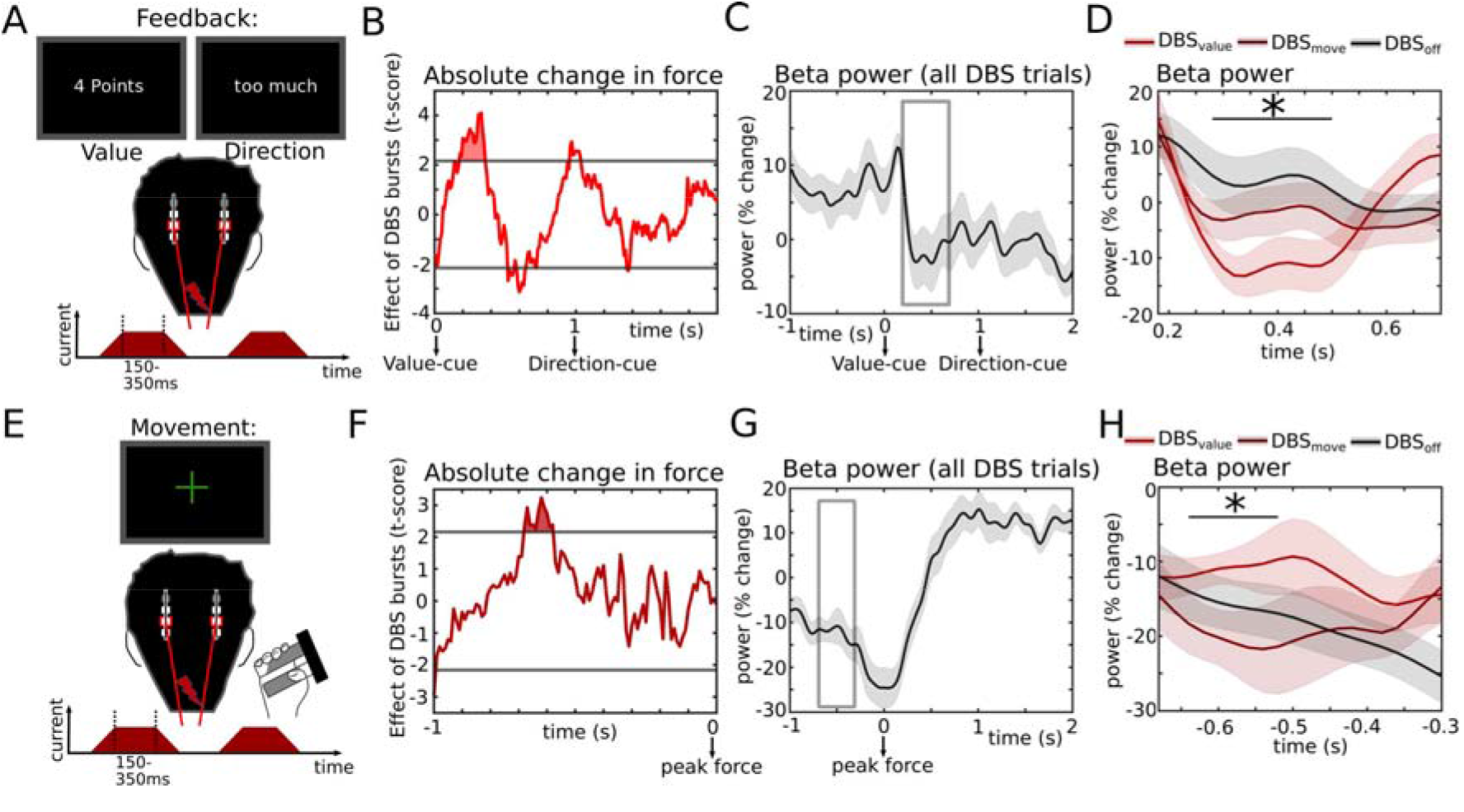
Causal effects of STN activity on action adaptation **A.** Bursts of STN stimulation were applied at random time points throughout the task including the feedback time period (see also figure 1A). **B.** STN stimulation increased the absolute change in force on the next trial when applied in a time window from 180 to 340 ms (termed DBS_value_) after the Value-cue. **C.** Beta power from the stimulation session. The grey rectangle indicates the time window from which beta activity was extracted for evaluating effects of stimulation on beta power. **D.** Stimulation at DBS_value_ reduced beta power compared to control trials in which DBS was applied in a time window from 680 to 580 ms before peak force (termed DBS_move_). Beta power from the no-stimulation session (DBS_off_) is also plotted. The black line with a ‘*’ indicates the time window with a significant difference between DBS_value_ and DBS_move_. **E.** Same as A for time windows aligned to the movement. **F.** STN stimulation increased the absolute change in force on the current trial when applied in a time window from 680 to 580 ms (DBS_move_) before peak force. **G.** Same as C for time windows aligned to the movement. **H.** DBS_move_ reduced beta power compared to control trials (DBS_value_). Beta power from the no-stimulation session (DBS_off_) is also plotted. The black line with a ‘*’ indicates the time window with a significant difference between DBS_move_ and DBS_value_. In B and F horizonal grey lines show the cluster-building threshold and filled areas indicate significant clusters. Shaded areas in C, D, G and H represent S.E.M. DBS, deep brain stimulation.

In a final analysis, we then aligned data to the movement and conducted the equivalent analysis as above (figure 3E). This revealed a significant, positive effect of STN stimulation on absolute change in force in a distinct time window from 680 to 580 ms before peak force (P_cluster_ < 0.05, figure 3F), hereafter termed DBS_move_. Here, stimulation led to a stronger absolute change in force on the current trial, but did not affect whether the force increased vs. decreased (suppl. figure 3D). Of note, DBS_move_ was in close proximity to the period in which STN beta power reflected absolute change in force off stimulation (440-300 ms before peak force, see figure 2E). Analogously to the analysis of DBS_value_, we therefore extracted beta power from the window comprising these periods (i.e. from 680 to 300 ms before peak force, see grey rectangle in figure 3G) now comparing DBS_move_ to DBS_value_ as control trials. This revealed a reduction of beta power by stimulation during DBS_move_ up to 340 ms before peak force which reached significance from 640 to 520 ms before peak force (P_cluster_ < 0.05, figure 3H). Thus, here stimulation led to a stronger absolute change in force and reduced STN beta power prior to reaching peak force, where lower beta power normally (i.e. off stimulation) reflected this behavioral adaptation.

## Discussion

Among the many possibilities afforded by our environment which actions should we choose and how should we perform them? One solution to this problem is to learn the value of different actions through trial-and-error and compare their outcomes to an expected value. Outcomes surpassing the expected value should be reinforced, while poor outcomes require adaptation. This reinforcement learning framework has proven to be highly useful in predicting behavior and even appears to have direct correlates in cortex – basal ganglia networks ^5-9^. At better-than-expected outcomes striatal dopamine is released from dopaminergic midbrain neurons, which can increase cortical excitability and shape neural dynamics through enhancement of the net-excitatory direct basal ganglia pathway ^3, 10-14^. However, the mechanisms mediating action adaptation after poor outcomes are less well understood.

Recording activity directly from the STN, a central part of the net-inhibitory indirect pathway ^11^, we found that reduced levels of beta power were related to larger absolute deviations between actual and target force as well as between larger absolute adjustments of force irrespective of direction. That is, this relationship did not depend on whether the force had been too low or too high or whether it increased or decreased on the next trial. This is compatible with previous studies suggesting that STN is critically involved in *changes* of actions or motor states rather than merely kinematics ^19, 20, 28^. Another way to put this is that reduction of STN beta activity may reflect a control effort necessary for changing between neural and / or behavioral states. This control cost can be disambiguated from mere metabolic costs since it e.g. increases when an isometric contraction is reduced or terminated ^19^, or a weaker effector is used ^17^.

Combining STN stimulation and simultaneous recordings of STN activity we found that short bursts of STN DBS were sufficient to modify adaptive behavior when applied in critical time windows and their effects on STN beta activity were consistent with their effect on behavior. Reduction of STN beta activity within or close to the time windows where lower levels of beta normally (off stimulation) warranted change led to stronger force adaptation. This provides further evidence for the usefulness of STN beta activity as a read-out or feedback signal for adaptive DBS approaches.

Some studies have observed a temporary dip in dopaminergic firing after worse-than-expected outcomes ^8, 9^. Since it has been proposed that dopamine release is related to reductions in beta activity of the STN ^29^ this would predict that lower action values in our study should lead to increases in beta activity. However, we observed the opposite namely a decrease in beta activity after worse outcomes and stronger action adaptation in line with previous studies ^20, 21^ suggesting a more intricate relationship between dopamine release and beta power ^30^. It should also be noted that STN beta activity mainly localizes to subparts of the STN connected to cortical motor areas rather than ‘reward’-related networks, which might provide distinct computations ^31-33^.

We also observed an increase in alpha activity after the Value-feedback, which however was not correlated to Value, i.e., it increased irrespective of negative or positive outcomes. Alpha activity in STN has mainly been related to attentional mechanisms, since it increases after salient stimuli ^34^ and is at rest coherent with temporo-parietal cortex ^35-37^ suggesting that it reflects a response to a salient cue rather than its content. In addition, we found a strong increase in gamma activity, which was maximal around the peak force. While gamma activity is closely related to force levels for stronger forces ^15, 16, 34^ and movement velocity ^34, 38, 39^ it appears not to be related to action adaptation, at least in the current paradigm. While these findings demonstrate the relative specificity of a component of STN beta activity as a signal reflecting action adaptation, we do not claim that complex behavior like action adaptation can be reduced to one simple LFP signal but will be reflected by multidimensional population dynamics across neural networks ^40^.

Are the mechanisms that we studied here relevant in the healthy state? We tested PD patients who express abnormally slow movements (especially during large amplitude movements and large forces ^41-43^), in particular OFF medication. To address this, we carefully designed the task using relatively low forces and limited cognitive demands, and assessed all patients in their ON medication state and immediately after DBS surgery when clinical impairment is less pronounced. Despite this, patients had difficulties in precisely producing lower forces, which might be related to impaired dexterity ^44, 45^ or the necessity to grip the dynamometer with a certain baseline force due to tremor (see supplementary text). However, patients had overall similar kinetics and, more importantly, similar measures of force adaptation as healthy people, which was the main interest of this study.

Another issue with the present investigation is that PD patients have reduced levels of dopamine and exaggerated STN beta activity when OFF medication ^29^, which might limit the generalizability of STN activity modulations during the task. As detailed above, all patients were studied ON medication and immediately after DBS surgery to mitigate this effect. Furthermore, there is a vast literature beyond STN LFP recordings in PD patients demonstrating neural correlates of force production in the basal ganglia including non-invasive recordings with functional magnetic resonance imaging in healthy people ^46, 47^, invasive electrophysiological recordings in healthy non-human primates ^48, 49^ and invasive electrophysiological recordings in humans with neurological disorders other than PD ^50^. In addition, cortical beta oscillations have been shown to reflect action adaptation in healthy humans using electroencephalography ^51, 52^. Together, this suggests that our findings may generalize beyond the studied patient group.

In summary, we here demonstrate that action evaluation and adaptation are reflected by dynamic STN beta activity and that causal manipulation of such activity can modify action adaptation in humans. More broadly, our results are in line with dynamic reductions in beta activity underlying adaptive processing. Future studies are warranted to assess whether this can be leveraged to re-establish physiological processing and assist motor execution and even learning ^53^ in patients suffering from neurological disorders.

## Methods

### Participants

We recruited sixteen patients with Parkinson’s disease (PD), who had undergone STN DBS surgery prior to the experimental recordings at University Medical Center at the Johannes Gutenberg University Mainz, Germany. Clinical details are listed in supplementary table 1. Lead localization was verified by monitoring the clinical effect and side effects during operation, as well as through postoperative stereotactic computerized topography (CT), see supplementary figure 6. Bilateral STN LFP were recorded and DBS applied through externalized electrode extension cables. The experiment was conducted in the immediate postoperative period 1-3 days after insertion of the DBS lead (Abbott 6170™), before implantation of the subcutaneous pulse generator, in the ON medication state. As a control group, we enrolled 16 healthy control (HC) participants without any neurological or psychiatric conditions. The groups did not differ regarding age (PD: 66 ± 13 years; HC: 67 ± 8 years, mean ± standard deviation; t_30_ = −0.227, P = 0.822, d = 0.08, independent samples t-test), handedness (1 left-handed person in each group as revealed by self-report, P = 1, Fischer’s exact test) or sex (14 male in PD group, 11 male in HC group, P = 0.394, Fischer’s exact test). All participants gave written informed consent to participate in the study, which was approved by the local ethics committee (State Medical Association of Rhineland-Palatinate) and conducted in accordance with the declaration of Helsinki. Two of the included PD patients (PD04 and PD07) did not participate in the second session with STN burst stimulation (see below) due to fatigue. One healthy participant had to be excluded due to miscalibration of the force device.

### Experimental task

We designed the experimental task so that participants had to constantly adapt the force they applied, whilst avoiding forces close to their maximum, or forces close to 0. Furthermore, the outcome should not be unambiguously predictable, so that participants had to wait for the feedback-cues after the movement, while also not being purely random. To this end, we computed trajectories of target force levels that noisily varied around their mean based on a decaying Gaussian random walk, which has been applied for studying decision-making under uncertainty ^24^. In particular the target force μ at each trial t was given by

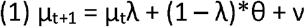

The start-value μ_1_ was drawn from a Gaussian distribution with a mean of 25 (% maximum voluntary contraction, MVC) and a standard deviation of 2, while ? was a noise term drawn from a zero-mean Gaussian with a standard deviation of 2. λ and θ were constants describing the rate of decay and the value towards which μ_t_ decayed to and were set to, respectively, 0.98 and 0.25 ^24^. The two trajectories (A and B, see figure 1B) used in this study were derived from simulations with the above given equations and kept constant across participants to facilitate comparisons between subjects. The resulting mean target force over trials was ∼20% MVC in both trajectories (i.e. slightly lower than θ). We used two trajectories for the target force levels, because patients participated in two sessions (one without stimulation and one with bursts of STN stimulation, see below). Since healthy participants only participated in one session (of note only the first session was compared between groups, i.e., PD patients off stimulation vs. HC) we counterbalanced the order of trajectories between session A and session B for PD patients and matched trajectories between patients and HC.

MVC was calculated as the median out of three attempts to press a manual dynamometer as hard as possible. For these attempts and all movements throughout the task, participants were instructed to apply relatively short force grips (in contrast to long-lasting isometric contractions) to avoid fatigue.

After estimating the MVC, participants performed a short training session in which they could get accustomed to the dynamometer and the initial force level. For this session the participants were told that the target force would be 25% of their maximal force, which was kept constant across the 10 training trials.

Next, the experimental session was conducted. The participants’ goal was to collect a maximum number of points by exerting forces that were as close to the target force level as possible at each trial. Participants were aware that the initial level was close to the training session, but that this would change over time. Since the target force was not shown on the screen, participants had to infer this based on the feedback they received. At the beginning of each trial a white fixation cross was shown for an average duration of 0.75 s (randomly jittered between 0.5 – 1s). When the fixation cross turned green, participants were instructed to produce the force they predicted to be as close as possible to the target force. The movement was allowed any time in a 2.5 s window and the fixation cross remained green for this whole duration irrespective of the timing of the movement. Participants were discouraged from exerting multiple presses even if they perceived that their first press was suboptimal, and to refrain from any movement after the first press and wait for the feedback. After this 2.5 s window a black screen was shown for 1-1.5 s after which the Value-feedback was presented for 1s. Values ranged from 0 (worst) to 10 points (best) and depended on the linear distance of the actual force from the target force, i.e., 0-1 % MVC difference resulted in 10 points, 1-2% MVC difference in 9 points, etc. Any difference > 10% MVC resulted in 0 points. After this Value-feedback, the Direction-feedback was shown for 1 s indicating whether the actual force had been “too much” (German: “zu viel”) or “too little” (German: “zu wenig”), after which the next trial began with a white fixation cross. The experimental session comprised 100 trials corresponding to ∼10 minutes. At the end of the session the sum of collected points was shown.

All cues were presented on a MacBook Pro (MacOS Mojave, version 10.14.6, 13.3 inch display, 60 Hz refresh rate) using PsychoPy v1.8 ^54^ implemented in Python 2. The display was viewed from a comfortable distance of ∼50 cm. Hand grip force was measured with a dynamometer (MIE Medical Research, Leeds, U.K.), which the participants held in their dominant hand with their forearm comfortably positioned on the armrest of the chair. Two people in each group used their non-dominant left hand, because of discomfort on the right side. The analogue force measurements were analogue-to-digital converted and sent to the PsychoPy software through a labjack u3 system (Labjack Corporation, Lakewood, CO, USA) as well as to the LFP recording device. In PsychoPy force was converted to % MVC for each individual participant. Task events were synchronized with the analogue force and LFP recordings (as well as DBS bursts in the second experiment, see below) by a TTL pulse that was sent from Psychopy to the recording software through the labjack system.

### Analysis of behavioral data

All trials without responses or with more than one response were excluded. The remaining trials were compared between groups across multiple measures of force production and adaptation to assess the generalizability of the results.

Force production: For each participant we calculated the mean peak force (peak force minus baseline), mean peak yank (first derivative of force), mean peak negative yank, and area under the curve (AUC, area between exerted force and baseline) and compared these variables between groups using independent samples t-tests. The baseline was computed as median of a 5s window centered on the Go-cue (i.e., 2.5 s before until 2.5 s after onset of the green fixation cross) at each trial to account for putative baseline drifts, which was visually inspected at each trial. We also compared the reaction time (mean time from Go-cue to peak force), the MVC (median of three attempts, see above) as well as the peak yank-to–peak force slope (Fisher z-transform of Pearson correlation coefficient) between groups. Force adaptation: Root mean squared error (RMSE), average Value-feedback (which is closely related to RMSE, since larger errors will results in fewer points collected), mean force error (mean difference in actual vs. target force, which reflects whether on average too much or too little force was applied), mean actual force at low vs. high target force levels (after median split of target force levels), coefficient of variation (CV, standard deviation of force divided by mean force), and mean by-trial absolute change in force (how much did participants on average change their force from trial to trial) were calculated and compared between groups using independent samples t-tests. To assess whether participants in general were able to follow task instructions we also calculated Pearson correlations between actual and target force (successful performance would predict a positive correlation) as well as Value-feedback and absolute change in force on the next trial (successful performance would predict a negative correlation). For all measures that reflected between-trial adaptation only trials where the previous trial also had been valid (i.e., two consecutive valid trials) were included. Throughout the analysis all data were tested for normality using Lilliefors test before conducting parametric tests. All results are listed in supplementary table 2.

### Processing of STN LFPs

LFPs were sampled from bilateral STN at 2048 Hz, bandpass filtered between 0.5 and 500 Hz and amplified with a TMSi porti device (TMS International, Enschede, The Netherlands). The same system was used for recording the force measures and TTL pulses (see above) through auxiliary input channels. The whole recording was visually inspected for artifacts off-line in Spike2 (Cambridge Electronic Design, Cambridge, UK) and noisy trials were rejected. After artifact rejection (on behavioral and neurophysiological grounds) ∼78 trials per patient and 1246 trials in total remained. Further analysis of the data was performed using FieldTrip ^55^ implemented in Matlab (R 2019a, The MathWorks, Natick, MA, USA). All scripts will be made available on https://data.mrc.ox.ac.uk. The data were imported to Matlab, high-pass filtered at 1 Hz using a 4^th^ order Butterworth filter, bandstop filtered between 49-51 Hz (FieldTrip function *ft_preprocessing*), and downsampled to 200 Hz using an anti-aliasing filter at 100 Hz (*ft_resample*). A bipolar montage was created from the monopolar recordings by computing the difference between the most dorsal omnidirectional contact and the neighboring three dorsal directional contacts, between the three dorsal and corresponding three ventral directional contacts, as well as the three ventral directional contacts and neighboring most ventral omnidirectional contact resulting in 8 bipolar channels per STN (*ft_apply_montage*). For each bipolar channel the data were transformed to the frequency domain using the continuous Morlet wavelet transform (width=7, *ft_freqanalysis*) for frequencies from 2 to 100 Hz using steps of 1 Hz and 20 ms throughout the whole recording. Power of each frequency was baseline corrected (*ft_freqbaseline*) relative to the mean power of that frequency across the whole recording ^56^ excluding time periods with large artefacts. The resulting spectra were epoched and aligned with, respectively, peak force and feedback-cues (*ft_redefinetrial*). In order to identify the bipolar contact, which showed the strongest task-related modulation, we analyzed all contacts with respect to their changes in movement-related beta (frequency of maximal modulation defined individually between 13-30 Hz) activity. We chose this as a functional localizer, because STN beta activity is localized within the dorsal STN ^57, 58^ and movement-related beta power modulation was present (defined as >15% reduction during movement) in all hemispheres. For each hemisphere the contact with the strongest movement-related decrease in beta power was chosen and these contacts then averaged across hemispheres resulting in one STN channel per patient. We confirmed the validity of this functional localizer approach by conducting lead localization analysis (see below).

### Electrode localization

Electrode localization was carried out using the Lead-DBS toolbox (v.2.5.2; https://www.lead-dbs.org/) with default parameters as described elsewhere ^59^. Briefly, using Advanced Normalization Tools (ANTs) preoperative magnetic resonance imaging and postoperative CT scans were corrected for low-frequency intensity non-uniformity with the N4Bias-Field-Correction algorithm, co-registered using a linear transform and normalized into Montreal Neurological Institute (MNI) space (2009b non-linear asymmetric). Brain shifts in postoperative acquisitions were corrected by applying the “subcortical refine” setting as implemented in Lead-DBS ^60^. The reconstructed electrodes (marked at contacts, which were used for LFP recordings and stimulation) were then overlaid on the STN to confirm proper targeting, see supplementary figure 6. Imaging data was not available in 2 patients.

### Statistical analysis of STN LFPs

Based on previous studies ^15-18^ we had a clear a-priori hypothesis about the spectral characteristics of STN activity relevant for force adaptations, namely the beta-band (∼13-30 Hz). To assess whether STN beta power in the current study was relevant for action evaluation and adaptation we applied the following analyses:

i) Feedback period: We aligned changes in STN beta power to the feedback cues (the Direction-cue was shown at a fixed interval of 1s after the Value-cue) and applied a linear mixed-effects (LME) model (Matlab function *fitlme*) using single trial beta power as dependent variable and Value-feedback as well as Direction-feedback as predictors.

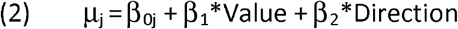

While the intersect was allowed to vary between each participant *j* (random effect) the slopes were fixed effects. All single trial values of STN beta power were z-scored by subtracting the mean and dividing by the standard deviation for each patient. Trials with z-scores > 3 were excluded (<1% of trials). Value-feedback (shown at the first feedback-cue) ranged from 0 (worst) to 10 (best). Direction-feedback (second feedback-cue, i.e., too little vs. too much force) was calculated as (10-value) multiplied by −1 (too little) or +1 (too much) resulting in a scalar ranging from −10 (force was > 10% MVC lower than target force) to +10 (force was > 10% MVC higher than target force), while 0 indicates no error.

We conducted these LME for 100 ms long moving windows of mean STN beta power, which were shifted by 10 ms from 0 (onset of Value-feedback) to 2 s (onset of Direction-feedback was at 1 s). The resulting t-values were then plotted over time, thresholded (corresponding to p<0.05) and the resulting clusters, which consisted of all time points that exceeded the initial threshold, were compared against the probability of clusters occurring by chance by randomly shuffling the trial order of STN beta power using 1000 permutations ^61, 62^. Of note, single trial beta power was shuffled across trials, while the order of time windows within each trial was preserved. Only clusters in the observed data that were larger than 95% of the distribution of clusters obtained in the permutation analysis were considered significant and marked as P_cluster_ < 0.05.

ii) Movement period: We aligned STN beta power to peak force and conducted LMEs as described for ‘Feedback period’ using predictors reflecting force adaptation. Predictors were the absolute change in force (i.e., how much the force was adapted from the previous to the current trial) and change in force (negative for a decrease in force and positive for an increase in force compared to the previous trial). For example, in a trial in which force was reduced by 5% MVC compared to the previous trial the absolute change in force was 5 and the change in force was −5.

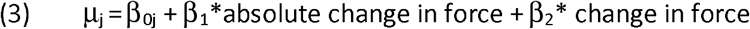

LMEs were conducted for 100 ms long time windows of mean STN beta power shifted by 10 ms from −1 s to peak force and corrected for multiple comparisons using cluster-based permutation tests as described above.

### Burst stimulation

After the first session patients had a short break of ∼30-60 min. During this the LFPs recorded from bilateral STN were processed and analyzed as described above, but instead of constructing bipolar channels from neighboring electrodes, two wider bipolar contacts were constructed to allow recording during stimulation of an intervening contact. First, directional contacts were averaged to form an omnidirectional contact (resulting in four omnidirectional contacts per STN). Then, a dorsal bipolar contact between the most dorsal and second most ventral contact and a ventral bipolar contact between the most ventral and the second most dorsal contact were created. This was done to compute the bipolar contact with the clearest movement modulation of beta activity, since this has been related to localization within or close to the dorsal STN ^13, 57^, and allows stimulation of the contact in between this bipolar pair to mitigate the stimulation artifact using common mode rejection ^63, 64^. The two bipolar contacts on each side were then compared regarding the extent of movement-related beta power modulation and the best contacts (i.e., with the clearest modulation) chosen as recording electrodes using the electrode in between as active contact for stimulation. DBS was applied using a custom-built device previously validated ^63^ in pseudo-monopolar mode using reference pads on the patients’ shoulders as anode. Frequency (130 Hz) and pulse width (60 µs) were fixed. To allow inference on timing-specific effects of stimulation DBS was applied in bursts. Mean DBS burst duration was 250 ms (drawn randomly from a uniform distribution between 150 and 350 ms) and mean burst interval was 150 ms (drawn randomly from a uniform distribution between 75 and 225 ms). These parameters were defined based on our previous study of closed-loop DBS ^63^ and were in simulations shown to result in DBS bursts occurring in ∼50% of trials in any given 100 ms time window during the experimental task allowing us to compare timing-specific behavioral effects of DBS vs. no-DBS. Stimulation was applied simultaneously to both hemispheres and ramped up and down to reduce paresthesia ^63, 64^. Ramp duration depended on the DBS intensity and ranged from 115 to 230 ms (see supplementary table 1). DBS intensity was titrated by slowly increasing the intensity of continuous DBS on each side and evaluating clinical effects on Parkinsonian symptoms as well as putative side effects by a trained clinician (DMH). When the threshold for clinical effects was reached the intensity was noted and, in case of side effects, slightly decreased. We evaluated this procedure by performing double-blind UPDRS-III scores (upper and lower limb bradykinesia, rigidity and tremor scores) in continuous DBS ON vs. OFF. This showed a consistent improvement in clinical scores on average from 27.4 to 20.2 (t_13_ = 6.151, P < 0.001, d = 1.64, paired samples t-test) confirming that the chosen intensities were clinically effective. We then used this intensity for burst stimulation while patients performed the same experimental task as described above. None of the patients reported paresthesia during the experiment.

### Effects of burst stimulation on force adaptation

Timing-specific effects of STN burst stimulation were analyzed using a moving-window approach ^63^. Stimulation intensity at each sample was saved in the recording software and imported to Matlab along with the TTL pulse (signaling the response), downsampled to 1000 Hz and binarized (0 for no stimulation, 1 for stimulation). Since intensities during ramping up and down of stimulation were below the clinically effective intensity they were defined as no stimulation ^63^. For each trial, we noted for 100 ms long time windows if stimulation was applied or not (at any point during that window). This time window was shifted by 10 ms over 2000 ms (from 0 to +2000 ms) for the feedback-aligned data and over 1000 ms (from − 1000 ms to peak force) for the movement-aligned data. We also analyzed the percentage of trials in which stimulation was applied at any given time window, which confirmed that stimulation was applied between ∼40 and 50% of trials at all time windows (suppl. figure 3A & B).

Based on the findings from the LFP-regression analysis (see results) we hypothesized that stimulation would modulate the absolute change in force. To test this, for each time window we computed the mean absolute change in force for all trials in which stimulation was applied and all trials in which stimulation was not applied. At the second level, i.e. in the across-subjects analysis, we then tested whether this measure was affected by stimulation by performing cluster-based permutation tests ^61, 62^. At each time window we computed the effect of stimulation using a cluster-building threshold corresponding to p<0.05 and the resulting clusters, which consisted of all time points that exceeded the initial threshold, were compared against the probability of clusters occurring by chance by randomly shuffling between stimulation labels (stimulation versus no stimulation) of each participant using 1000 permutations. Only clusters in the observed data that were larger than 95% of the distribution of clusters obtained in the permutation analysis were considered significant and marked as P_cluster_ < 0.05. The time window in which DBS had significant behavioral effects after the Value-cue (see results) was termed DBS_value_, while the time window where DBS had significant behavioral effects aligned to the movement was termed DBS_move_.

As a control analysis, we repeated these analyses using change in force (rather than absolute change in force).

### Effects of burst stimulation on STN LFPs

Whilst stimulation was applied LFPs were continuously recorded through the two contacts neighboring the stimulation contact and a bipolar signal derived as previously described (i.e. wide bipolar recording). Despite common-mode rejection the artifact was clearly visible (see suppl. figure 4A) and its spectral characteristics were not strictly confined to the stimulation frequency and its harmonics (suppl. figure 4B & C). Hence the following artifact removal procedure was applied. The data were imported to Matlab, high-pass filtered at 4 Hz and low-pass filtered at 100 Hz using a 4^th^ order butterworth filter, demeaned and detrended (*ft_preprocessing*). After visual inspection of the LFPs from each patient a common threshold was set at 10 µV. This was chosen, because the remaining (i.e. after filtering) stimulation artifact, but not physiological LFPs (in the interval of stimulation bursts), consistently crossed this threshold. At each sample the signal was removed if it crossed the threshold (∼9% of the data) and replaced by linear interpolation of the neighboring non-noisy signals. Afterwards the data were downsampled to 200 Hz and the subsequent time-frequency analysis was identical to the LFP recordings described above. An example of single subject beta power is shown in suppl. figure 4D & E, subject-averaged spectra are shown in suppl. figure 4F & G.

After preprocessing and artifact correction, we first assessed the overall effect of stimulation on beta power by aligning beta power to onset of stimulation (after ramping) and normalizing it to the mean beta power when no stimulation was applied (i.e. during the stimulation interval). As expected ^25-27^, this showed a clear stimulation-related reduction in beta power, see suppl. figure 5.

Then, we analyzed the effects of timing-specific stimulation (during DBS_value_ and DBS_move_) on STN beta power. Since the LFP regression analysis showed a relationship between levels of STN beta power and behavioral changes (see results) and since stimulation reduced beta power (suppl. figure 5) we asked whether stimulation at these specific time points also affected beta power at specific windows. To this end, we first extracted beta power from the feedback-period in which stimulation affected behavior (DBS_value_) and where STN beta power usually (i.e. off stimulation) correlated with Value, i.e. from 180 to 700 ms after Value-feedback (grey rectangle in figure 3C). We then compared trials in which stimulation had been applied in the critical time window (DBS_value_) and compared this to trials in which stimulation also affected behavior but in a distinct window (DBS_move_). The rationale for this was to match the two conditions as well as possible regarding recording technique and signal quality (both conditions were from the stimulation session with wide bipolar contacts). We also plotted beta power from the off-stimulation session as reference. However, it should be noted that in these trials both recording technique (narrow bipolar) and signal quality (no stimulation artefacts) were different.

We conducted the analogous analysis for movement-aligned data comparing beta power of DBS_move_ and DBS_value_ in the time window from 680 to 300 ms before peak force. The statistical analyses were conducted using cluster-based permutation tests by shuffling between stimulation labels (DBS_move_ vs. DBS_value_) as described above.

## Acknowledgements

DMH is supported by a postdoctoral grant from the Independent Research Fund Denmark (0168-00014B). PB is supported by the Medical Research Council (MC_UU_00003/2). RB is supported by the Medical Research Council (MC_UU_00003/1).

## Data availability

Original data are available upon reasonable request to the corresponding author. All scripts will be made freely available on https://data.mrc.ox.ac.uk.

