## supplemental text, figures and tables for "Neural underpinnings of action adaptation in the subthalamic nucleus"

Supplementary information:

Supplementary text:

Patients showed overall poorer task performance compared to healthy controls regarding the difference between actual and target force resulting in lower average Value-feedback (t_29_ = 3.416, d = 1.23, P = 0.002, see figure 1C and suppl. table 2 for measure of force adaptation). This was mainly due to patients having difficulties in producing very low forces (force trajectories are shown in suppl. figure 1). Thus, patients produced significantly higher forces compared to HC during low target force trials, while there were no differences between groups in high target force trials (for statistics see suppl. table 2). However, these difficulties did not impact patients’ overall ability for force adaptation as indicated by similar force level variability (coefficient of variation, t_29_ = 0.572, d = 0.21, P = 0.572) and mean by-trial absolute change in force (t_29_ = 0.346, d = 0.12, P = 0.732) between groups (single participant measures shown in figure 4D). Importantly, whilst on average overshooting force, patients were still able to reduce their force even when target force levels were low (correlation between force overshoot and decrease in force on the next trial during low target force trials in PD patients: mean rho = 0.364, P < 0.001). This suggests that the group difference in average Value-feedback was due to patients having difficulties in producing very low forces presumably related to impaired dexterity or tremor (an exploratory Pearson correlation between individual tremor scores and force overshoot showed a correlation coefficient of 0.51, P = 0.052) rather than a deficit in force adaptation.

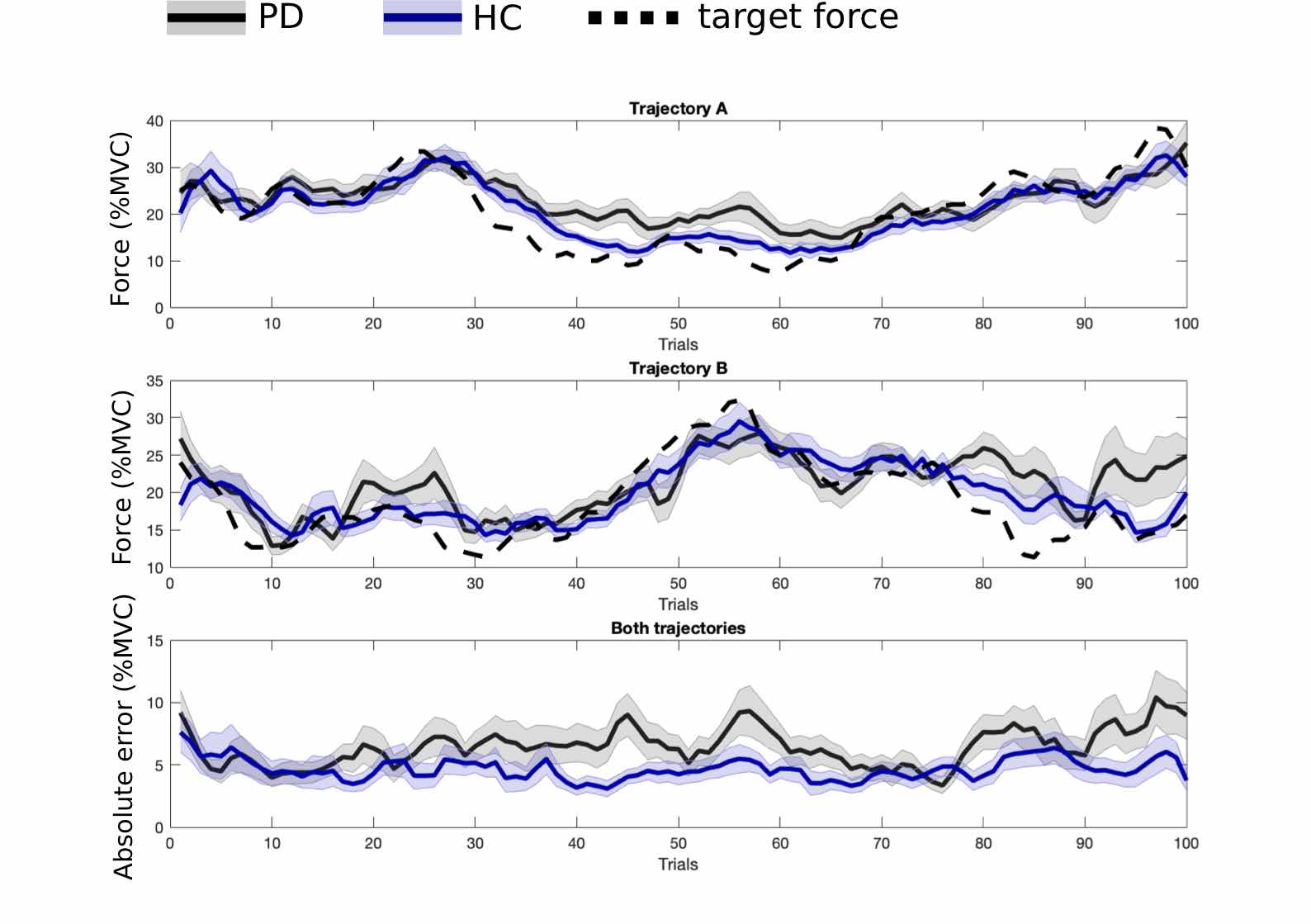

**Supplementary figure 1**: Actual vs. target force. Mean actual force (black: Parkinson’s disease (PD) patients; blue: healthy controls (HC)) are plotted along with target force (dotted lines) for all trials of trajectory A (upper panel) and trajectory B (middle panel). Participants showed the strongest errors when the target force was very low (e.g. middle trials of trajectory A or last trials of trajectory B). This was most pronounced in PD patients as can be seen in the lower panel showing the absolute error over trials. MVC, maximum voluntary contraction. Shaded areas represent S.E.M.

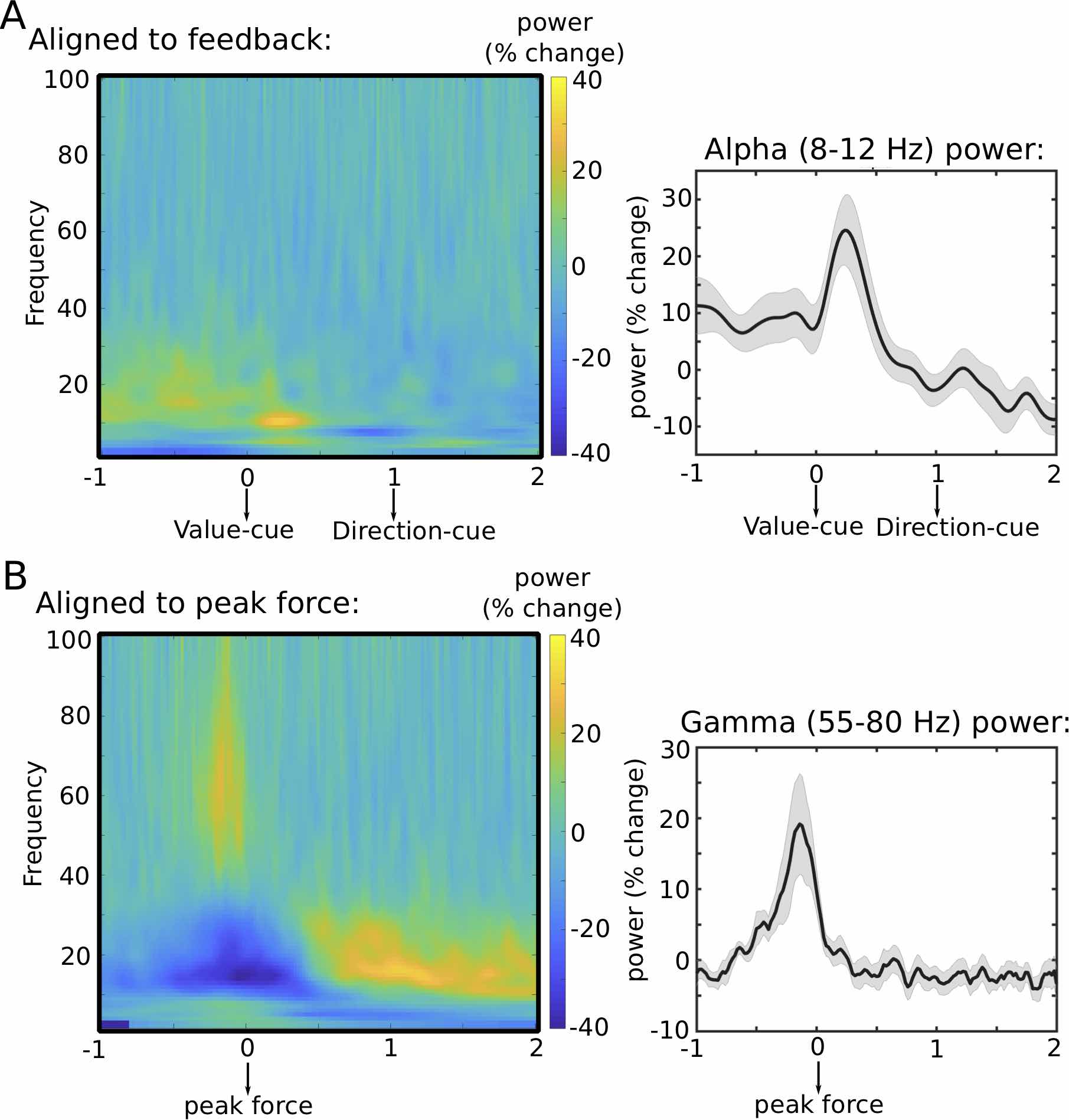

**Supplementary figure 2.** Time-frequency spectra. **A.** Spectra aligned to the feedback showing a strong increase in STN alpha (8-12 Hz) power from ~0-0.5 s after the Value-feedback. The right panel shows mean alpha power aligned to the feedback. **B.** Spectra aligned to peak force showing a strong increase in STN gamma (55-80 Hz) power from ~0.3 before peak force until peak force. The right panel shows mean gamma power aligned to peak force. Shaded areas in the right panels represent S.E.M.

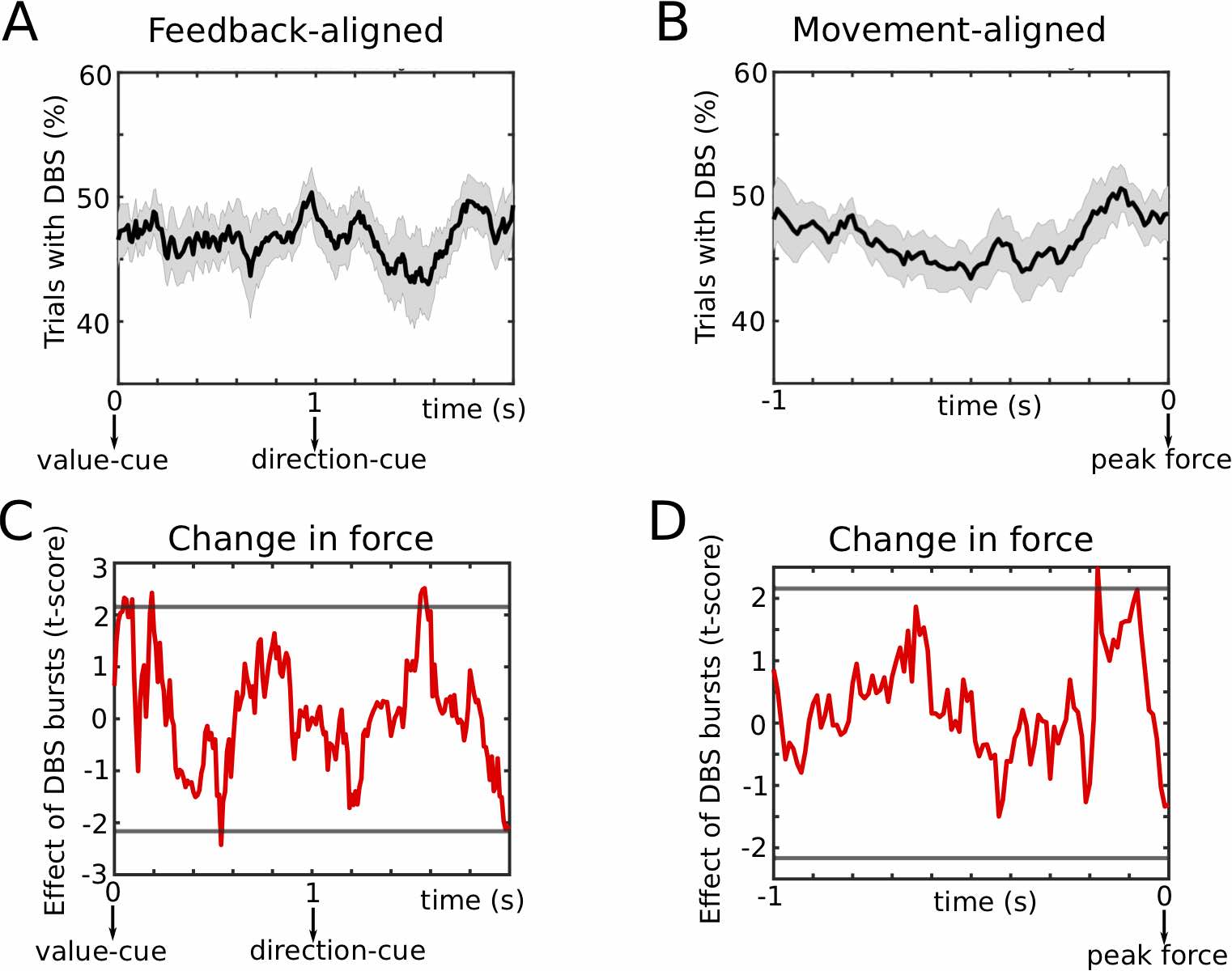

**Supplementary figure 3.** Burst stimulation. **A.** Stimulation occurred on ~50% of trials for any given 100 ms window throughout the feedback time period. **B.** Same as A for peak force aligned data. **C.** There were no significant effects of stimulation on change in force (positive values for increase in force, negative values for decrease in force) on the next trial when aligning data to the feedback cues. **D.** Same as C for peak force aligned data. Shaded areas in A and B represent S.E.M. Horizontal grey lines in C and D show the cluster-building threshold.

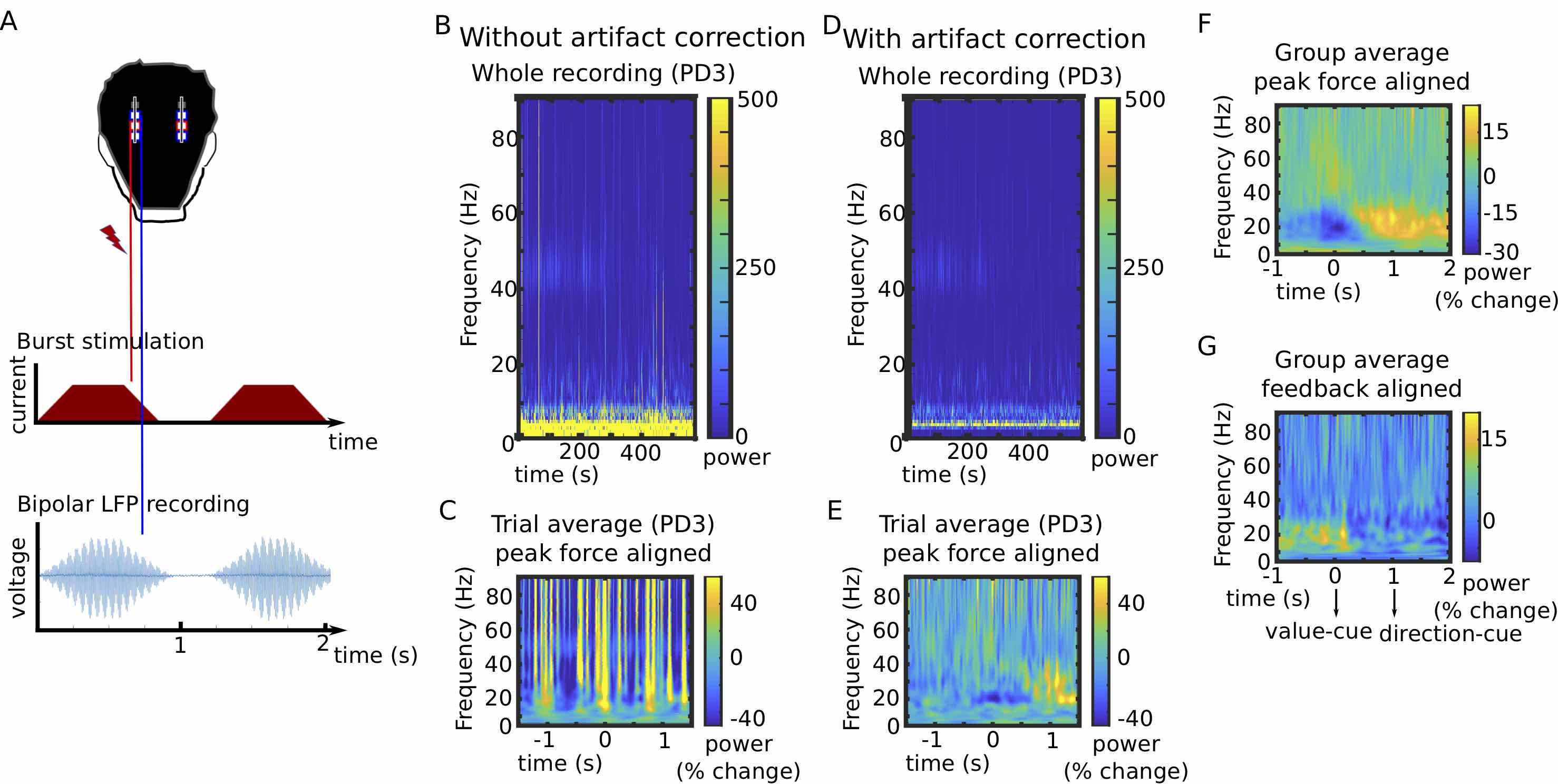

**Supplementary figure 4.** Stimulation-induced artifact. **A**. During the stimulation session, local field potentials were recorded from bipolar contacts surrounding the stimulation electrode. Despite common mode rejection the artifact was clearly visible in the un-processed data. **B.** Example of time-frequency spectrum without artifact correction (see methods for details regarding artifact correction) for an example patient (PD03). The spectral properties of stimulation-related artefacts were not restricted to the stimulation frequency and its harmonics. **C.** When the artefacts were strongly expressed, as in this patient, they obliterated the normal movement-related beta power modulation in the trial averaged data (peak force aligned). **D.** Same as B but after artifact correction**. E.** After artifact correction the normal (i.e. as observed in the off stimulation session) beta modulation can be seen in the trial averaged data (peak force aligned). **F**. Group average of peak force aligned spectrum after artifact correction (compare to supplementary figure 2B). **G**. Group average of feedback aligned spectrum after artifact correction (compare to supplementary figure 2A). Beta power time series extracted from F and G are also shown in figure 3C and 3G.

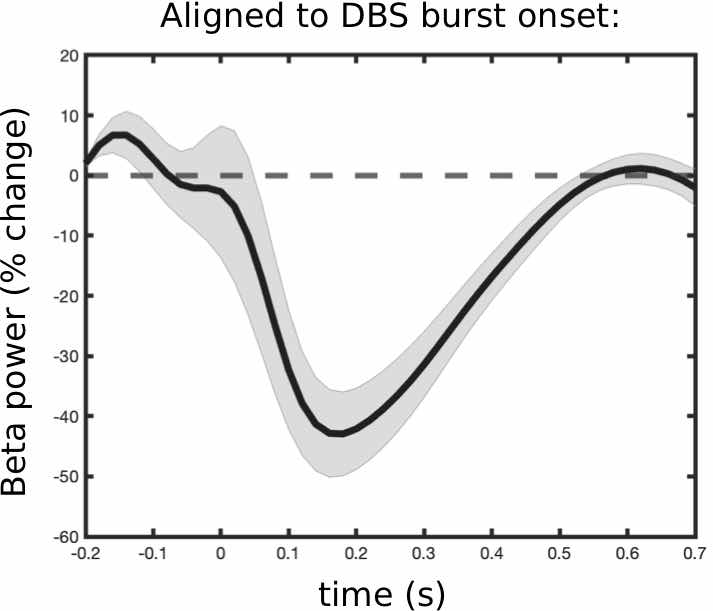

**Supplementary figure 5.** Effect of stimulation bursts on beta power. Beta power is aligned to onset of stimulation (after ramping) and normalized to the time period where no stimulation was applied. Stimulation led to a ~40% decrease in beta power, which returned to baseline after ~0.5 s (mean burst duration was 250 ms +/- 100 drawn from a uniform distribution). Shaded areas represent S.E.M.

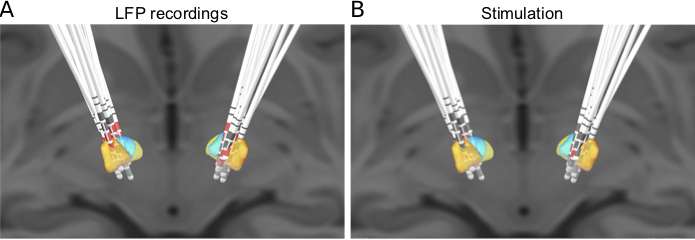

**Supplementary figure 6.** Electrode localization. **A.** Reconstructed bilateral leads are overlaid on subthalamic nucleus (STN; orange: STN area mainly connected to ‘motor’ regions, blue: STN area mainly connected to ‘associative’ regions, yellow: STN area mainly connected to ‘limbic’ regions). Electrodes from which bipolar local field potential (LFP) signals were analyzed are marked in red. **B.** Same as A, but marked electrodes indicate contacts which were used for burst stimulation.

| # | Age & gender | UPDRS-III OFF/ON levodopa | UPDRS-III limb OFF/ON DBS | Disease duration | Main symptom | Reason for surgery | Medication (LEDD) | DBS parameters Left / Right |
| --- | --- | --- | --- | --- | --- | --- | --- | --- |
| 1 | 60 male | 33 / 14 | 22 / 17 | 6 | Bradykinesia | ON-OFF fluctuations | 1569 mg | D-D, 1.7 mA, 0.215 s |
| 2 | 69 female | 25 / 17 | 24 / 20 | 7 | Bradykinesia | Wearing OFF | 1285 mg | D-V, 2.4 mA, 0.184 s |
| 3 | 62 male | 32 / 24 | 10 / 4 | 5 | Bradykinesia and tremor | Dyskinesia | 400 mg | D-D, 2.2 mA, 0.169 s |
| 4 | 79 male | 28 / 12 | n/a | 16 | Bradykinesia | ON-OFF fluctuations | 1170 mg | n/a |
| 5 | 68 male | 59 / 50 | 32 / 29 | 13 | Bradykinesia | ON-OFF fluctuations | 1448 mg | D-D, 2.6 mA, 0.200 s |
| 6 | 70 male | 46 / 31 | 23 / 13 | 13 | Bradykinesia | Dyskinesia | 1548 mg | V-V, 1.3 mA, 0.200 s |
| 7 | 67 male | n/a | n/a | 19 | Tremor | Wearing OFF | 1480 mg | n/a |
| 8 | 78 male | 44 / 20 | 23 / 13 | 16 | Bradykinesia | ON-OFF fluctuations | 1000 mg | D-D, 2.50mA, 0.153 s |
| 9 | 71 male | 34 / 32 | 27 / 21 | 4 | Tremor | Tremor | 450 mg | D-D, 2.0 mA, 0.153 s |
| 10 | 68 male | 63 / 33 | 37 / 31 | 30 | Bradykinesia | ON-OFF fluctuations | 892 mg | D-D, 1.1 mA, 0.169 s |
| 11 | 73 female | 45 / 26 | 27 / 27 | 11 | Tremor | Tremor | 900 mg | V-V, 2.0 mA, 0.153 s |
| 12 | 73 male | 42 / 23 | 32 / 26 | 6 | Bradykinesia | Gait difficulties | 1514 mg | D-D, 2.5 mA, 0.192 s |
| 13 | 49 male | 24 / 9 | 36 / 20 | 10 | Bradykinesia | Dyskinesia | 455 mg | D-D, 2.0 mA, 0.153 s |
| 14 | 80 male | 62 / 54 | 39 / 27 | 15 | Tremor | Tremor | 705 mg | D-D, 3.0 mA, 0.230 s |
| 15 | 66 male | 23 / 18 | 16 / 13 | 17 | Bradykinesia | ON-OFF fluctuations | 1863 mg | D-D, 1.5 mA, 0.115 s |
| 16 | 25 male | 54 / 32 | 36 / 32 | 3 | Bradykinesia | Dyskinesia | 1065 mg | D-D, 1.6 mA, 0.123 s |

**Supplementary table 1.** Age and disease duration are given in years. Clinical scores are given as total score of the MDS Unified Parkinson’s disease rating scale (UPDRS) part III for levodopa ON/OFF and as items 3-8 & 14-18 (limb scores) for DBS ON/OFF. Medication is given in levodopa-equivalent daily dose (LEDD). All patients received levodopa, 12 patients received a dopamine-agonist, 10 patients a catechol-O-methyltransferase (COMT) inhibitor, 9 patients a monoamine-oxidase (MAO) inhibitor and 4 patients amantadine. D and V indicate whether the respectively more dorsal or ventral contact was chosen as active contact in the left and right hemisphere, after which DBS intensity is given in mA and ramp time is given in seconds. n/a, not available.

|  | Mean ± standard deviation (PD vs. HC) | t_dof_-value,p-value, Cohen’s d |
| --- | --- | --- |
| Force production: |  |  |
| MVC (N) | 182.3 ± 58 vs. 211.2 ± 60 | t_29_=-1.366, P=0.171, d=0.49 |
| Mean peak force (N) | 22.5 ± 10 vs. 22.0 ± 7 | t_29_= -0.182, P=0.857, d=0.07 |
| Mean peak yank (N/dt) | 0.20 ± 0.1 vs. 0.17 ± 0.1 | t_29_=-0.949, P=0.351, d=0.34 |
| Mean peak negative yank (N/dt) | 0.23 ± 0.1 vs. 0.22 ± 0.1 | t_29_=-0.180, P=0.859, d=0.06 |
| AUC (N*ms) | 8780 ± 6236 vs. 7884 ± 4126 | t_29_=-0.469, P=0.643, d=0.17 |
| Time from Go-cue to peak force (s) | 1.13 ± 0.4 vs. 1.05 ± 0.2 | t_29_=-0.645, P=0.524, d=0.23 |
| Peak force–to-peak yank slope | 0.09 ± 0.04 vs. 0.07 ± 0.02 | t_29_=-1.484, P=0.149, d=0.53 |
| Force adaptation: |  |  |
| RMSE (%MVC) | 8.5 ± 2 vs. 6.1 ± 1 | **t_29_=-3.371,** **P=0.002, d=1.21** |
| Average points (au) | 4.9 ± 1 vs. 6.1 ± 1 | **t_29_=3.416,** **P=0.002, d=1.23** |
| Mean force difference (%MVC) | 2.5 ± 3 vs. 0.7 ± 1 | **t_29_=-2.454,** **P=0.020, d=0.88** |
| Actual force during high target force* | 24.7 ± 3 vs. 24.6 ± 2 | t_29_=-0.069, P=0.945, d=0.02 |
| Actual force during low target force* | 20.1 ± 4 vs. 16.6 ± 1 | **t_29_=-3.487, P=0.002, d=1.25** |
| CV (unitless) | 0.33 ± 0.1 vs. 0.35 ± 0.1 | t_29_=0.572, P=0.572, d=0.21 |
| Mean by-trial absolute change in force (%MVC) | 5.3 ± 2 vs. 5.5 ± 2 | t_29_=0.346, P=0.732, d=0.12 |

**Supplementary table 2.** Overview of group comparisons. au, arbitrary units; AUC, area under the curve; CV, coefficient of variation; dof, degrees of freedom; dt, time derivative; ms, millisecond; MVC, maximum voluntary contraction; N, Newton; RMSE, root mean squared error; s, second. * These two effects were significantly different from each other when directly comparing them (t_29_=-2.661, d=0.956, P=0.013).
